# Emergence of travelling wave patterns in resource-mediated tissue competition

**DOI:** 10.64898/2026.08.11.744236

**Authors:** N. Briñas–Pascual, T. Alarcón, J. Calvo, P. Guerrero, R. Oliver–Bonafoux

**Affiliations:** Universidad Carlos III de Madrid, Departamento de Matemáticas Avda. Universidad, 30, Leganés, 28911, Spain; Grupo Interdisciplinar de Sistemas Complejos (GISC); Institució Catalana de Recerca i Estudis Avançats Passeig Lluís Companys, 12, Barcelona, 08010, Spain; Centre de Recerca Matemàtica Edifici C, Campus de Bellaterra, Cerdanyola del Vallès, 08192, Spain; Departament de Matemàtiques, Universitat Autònoma de Barcelona Edifici C, Campus de Bellaterra, Cerdanyola del Vallès, 08192, Spain; Barcelona Collaboratorium for Predictive and Theoretical Biology Wellington, 30, Barcelona, 08005, Spain; Departamento de Matemática Aplicada and Research Unit “Modeling Nature” (MNat), Universidad de Granada Avda. Fuentenueva s/n, Facultad de Ciencias, Granada, 18071, Spain; Dipartimento di Informatica, Università di Verona Strada Le Grazie 15, Verona, 37134, Italy

**Keywords:** Travelling waves, Reaction-diffusion equations, Tissue competition, Quasi-steady-state approximation, Phenotypic heterogeneity

## Abstract

The study of tissue dynamics has been stimulated during the last decades thanks to the use of quantitative descriptions, with the development of several theoretical and computational frameworks, many of them revolving around the notion of reaction-diffusion systems, eventually with additional structure variables beyond time and space. The use of structure variables can accommodate phenotypic traits. In this work, we study a family of competition models, where a given population depends on a resource (e.g. oxygen) and several populations are competing for it. Our quantitative description incorporates phenotypic traits and heterogeneity at the level of cell cycle variations, which influence replication rates via oxygen consumption. This enables us to replicate the fitness of specific subpopulations to environmental conditions (e.g. oxygen shortage or external influences). Using numerical simulations, we show that such models display dynamical pattern formation in the form of coupled travelling wave profiles that expand or retreat at the same wave speed. The full theoretical analysis of such dynamics is quite involved; to circumvent this difficulty, we introduce a quasi-stationary approximation for the resource dynamics. We find that this approximation can reproduce the overall behaviour very accurately, with the additional benefit of allowing theoretical treatment of the reduced model. In this way, we provide estimates on the wave speed which are numerically shown to be robust across a wide range of macroscopic parameters of the full model. The wave speeds are thus found to depend strongly on the proliferation rate of the fittest population, resembling a winner-takes-all dynamics.

## 1. Introduction

The dynamics of competing cell populations underlie some of the most consequential processes in biology, from the maintenance of healthy tissue homeostasis to the progressive expansion of tumour cell lineages into surrounding tissue Hanahan and Weinberg (2011); Marusyk et al. (2012). In these settings, individual cells do not act in isolation: they compete for space, signalling cues, and metabolic substrates. The outcome of that competition shapes tissue structure, function, and ultimately clinical prognosis. A feature that makes such systems particularly rich — and particularly difficult to understand — is phenotypic heterogeneity. Even within a nominally uniform tissue, cells may differ in their ability to progress through the replication cycle, to tolerate oxygen shortage, or to sustain proliferation under metabolic stress Chisholm et al. (2015); Ardaševa et al. (2020). These differences need not be genetically encoded; they can reflect cell-to-cell variation in the concentrations of regulatory molecules that govern the cell cycle de la Cruz et al. (2015); Bedessem and Stéphanou (2014), and they can be transient, context-dependent, and heritable over short timescales. Understanding how such heterogeneity shapes competition, and how it feeds back onto spatial organisation at the tissue scale, is a central challenge at the interface of quantitative biology and mathematical modelling.

A natural mechanism through which phenotypic differences become consequential is competition for a shared resource. When cell proliferation depends on resource availability — as is the case for oxygen in poorly vascularised or rapidly growing tissue — small differences in the efficiency with which subpopulations extract or utilise such resource produce differences in net growth rate. In a spatially extended system, this creates a self-amplifying feedback: a subpopulation with a marginally higher proliferation rate at a given resource level will consume the resource more rapidly locally, further suppressing the growth of its competitors Gatenby and Gawlinski (1996); Martínez-González et al. (2012). Even modest initial differences in cellular phenotype can therefore be amplified by the spatial dynamics into large-scale competitive asymmetries, potentially culminating in the progressive elimination of less fit subpopulations. This mechanism operates without recourse to genetic mutation or selection in the classical evolutionary sense, and it suggests that phenotypic heterogeneity alone — mediated by resource competition — may be sufficient to drive competitive takeover in tissue Chisholm et al. (2015); Ardaševa et al. (2020).

In spatially extended systems, the hallmark of such competitive dynamics is the emergence of travelling invasion patterns. Rather than two competing populations simply exchanging dominance uniformly across space, a coherent spatial structure develops: the fitter population advances as an invading front, while the less fit population retreats at the same speed, and the resource concentration reorganises accordingly. The importance of travelling wave solutions as a mathematical description of biological invasion was recognised in the foundational works of Fisher and Kolmogorov Fisher (1937); Kolmogorov et al. (1937), and their relevance to tumour growth and tissue competition has since been established in a broad class of reaction-diffusion models Sherratt (1993); Gatenby and Gawlinski (1996); Marchant et al. (2000, 2001); El-Hachem et al. (2021); Gallay and Mascia (2022); Colson et al. (2021); Lorenzi et al. (2025a). Analogous phenomena have been identified in phenotype-structured models incorporating chemotactic or haptotactic couplings Freingruber et al. (2025); Lorenzi et al. (2025b), as well as in stochastic and hybrid competition models de la Cruz et al. (2017). A recurring observation across these settings is that coupled wave structures emerge robustly after a transient, with the populations and resource all propagating at a common speed; the fittest population progressively colonises the domain while the less fit one retreats. The speed at which this process unfolds, and the conditions under which it occurs, carry direct biological meaning: they determine how quickly one cell lineage can displace another, and what phenotypic properties of the invading population are most consequential for the outcome.

To study these phenomena in a tractable yet biologically grounded setting, we introduce a family of coarse-grained reaction-diffusion models in which one or more cell populations compete for a single shared resource, here intended as a proxy for oxygen or any analogous rate-limiting metabolic substrate. Phenotypic heterogeneity is incorporated through an effective, resource-dependent proliferation rate, which propagates cell-to-cell variation in cell-cycle dynamics to the population level. Our model does this without resolving the full intracellular machinery. This coarse-graining is motivated by earlier multiscale and structured-population models de la Cruz et al. (2017, 2015); Bedessem and Stéphanou (2014), and it captures biologically meaningful growth rate differences between competing lineages — differences that arise, for instance, from variation in the rate at which cells traverse specific cell-cycle checkpoints under varying oxygen availability — while keeping the model analytically and numerically tractable. The resulting framework is deliberately general: the specific form of the proliferation function can be varied to reflect different biological contexts, and the qualitative behaviour we describe is robust to this choice.

The full coupled system, comprising population densities and resource concentration evolving together in space and time, is analytically untractable. A key simplification arises when the resource dynamics relax on a timescale that is fast relative to cell proliferation — a biologically reasonable approximation in many tissue settings, where oxygen diffuses and equilibrates far more rapidly than cells divide or migrate. Under this quasi-steady-state approximation (QSSA), the resource concentration at each point in space is slaved to the instantaneous local cell density, and the system reduces to a single equation per population with a nonlinear, density-dependent effective growth rate. This reduction is not merely a technical convenience: it clarifies the dominant feedback structure of the system, making explicit the way in which local cell density sets local resource availability, which in turn governs the local net proliferation rate. As our numerical results show, the QSSA provides a quantitatively accurate description of the full system’s behaviour across a wide range of biologically relevant parameters, and it makes the travelling wave problem accessible to rigorous analysis.

The present paper analyses travelling wave dynamics in both the single-population and two-population versions of these resource-mediated competition models. For the single-population model, the reduced system takes a form analogous to the classical Fisher–KPP equation Fisher (1937); Kolmogorov et al. (1937); Hadeler and Rothe (1975), and the analysis identifies the conditions under which a coupled population–resource travelling wave emerges together with an estimate of the invasion speed in terms of the proliferation rate at the uncrowded resource level. For the two-population competition model, the reduced system is a four-dimensional non-autonomous dynamical system not amenable to standard phase-plane methods. We exploit the fact that, under a suitable change of variables, it can be recast as a monotone cooperative system in the sense of Volpert et al. (1994), which yields existence of coupled travelling wave solutions with monotone profiles and a lower bound on the invasion speed. Our numerical simulations confirm that coupled travelling wave patterns emerge robustly across a wide range of parameter values, that the QSSA reduction is quantitatively accurate, and that the predicted invasion speed, governed principally by the proliferation properties of the fitter population evaluated at the critical resource threshold of the host, agrees well with the speed observed in simulations of the full model. Taken together, these results suggest that resource-mediated competition, even when phenotypic differences are modest, reliably produces a winner-takes-all spatial dynamic whose rate is set by the growth properties of the fitter lineage.

The paper is organised as follows. Section 2 introduces the single-population model, describes the non-dimensionalisation and numerical scheme, derives the QSSA, and analyses travelling waves for the reduced system. Section 3 extends the framework to the two-population competition model, establishes the existence of coupled travelling waves, and presents a systematic numerical assessment of the QSSA and the wave-speed prediction. Section 4 discusses the biological implications of the results and outlines directions for future work. Technical proofs are collected in the Supplementary Material.

### 1.1. Description of the model and results

Although our main interest lies in describing two competing tissues, it is instructive to consider first a simplified version of this situation, where we take into account just one cell population *n* = *n*(*t, x*), together with a given resource *c* = *c*(*t, x*). We take the unknowns to represent cell density and oxygen concentration (say), which evolve in time according to the following PDE system:

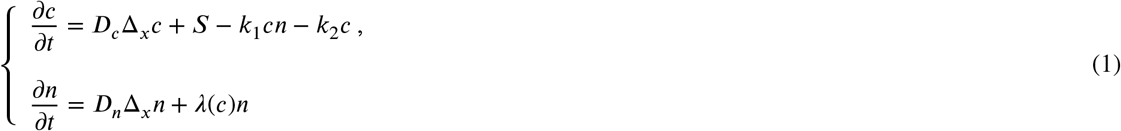

Here we consider that both species diffuse in space (***D***_*n*_ and ***D***_*c*_ represent the corresponding diffusion coefficients). We also consider that the population consumes the resource at a certain rate which is proportional to resource availability, therefore we describe the interaction between both species as a second order kinetics with rate *k*_1_. Moreover, we consider that the resource degrades, with rate *k*_2_, and also that there is a resource input, given by the source term *S*. Finally, *λ* = *λ*(*c*) is an effective proliferation rate (i.e. it can be negative, which reflects the fact that the members of the population can die), that depends on resource availability: the more resource is available, the faster the population will proliferate. In de la Cruz et al. (2017) this proliferation rate is given by a coarse-graining of a cell-cycle model; we explain this in more detail in Section 2 below. We can take the main properties verified in that representative situation to perform a more general analysis. During this document we shall thus take for granted the following *Assumptions*:

1. There exist numbers *d*_+_ ≥ 0 and *r*_−_ *<* 0 *< r*_+_ such that *λ* : (*d*_+_, +∞) ⟶ (*r*_−_, *r*_+_).
2. The effective proliferation rate *λ*(*c*) is strictly increasing as a function of the resource *c*.
3. There holds that *S > k*_2_*λ*^−1^(0).

As per Assumption 1, the proliferation rate can be either positive or negative (in which case it is interpreted as a net death rate for the population). It nevertheless increases when the resource availability does, thanks to Assumption 2. This assumption also guarantees that *λ*^−1^ : (*r*_−_, *r*_+_) ⟶ (*d*_+_, +∞) makes sense. In particular, *λ*^−1^(0) is well defined -this magnitude is the (positive) oxygen concentration yielding zero net proliferation rate. The remaining assumption is of a technical nature: item 3 is needed to ensure the existence of homogeneous equilibria -actually *S*/*k*_2_ would be the homogeneous equilibrium for the oxygen concentration in the absence of the population; we discuss this in more detail below.

The main features of this model are already met in spatial dimension one; in what follows, our investigation is restricted to this specific scenario. The additional phenomenology that is met for spatial dimension greater than one will be tackled elsewhere.

#### The competition model

Our main interest in this document are situations where several populations as described before compete for the same resource. We have in mind idealised scenarios in tumour prognosis where tumoural tissue proliferates against the surrounding healthy tissue. For those scenarios, a two-population model like the following one is representative:

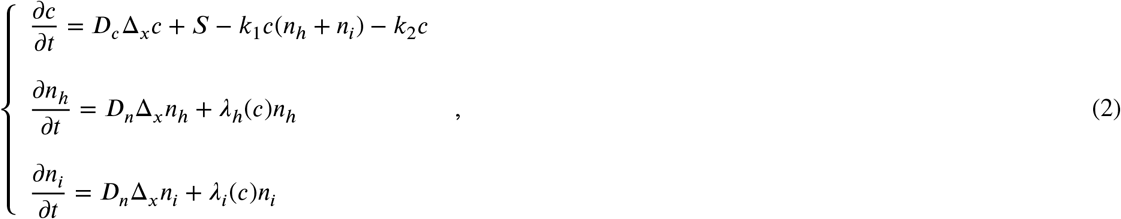

where the population *n*_*i*_ -”invader”-is fitter than *n*_*h*_ -”host”-, as the property *λ*_*i*_(*c*) *> λ*_*h*_(*c*), relating their respective proliferation rates, confers replicative advantage to the former. Such replicative advantage can have different causes; in the vein of de la Cruz et al. (2015, 2017), we can attribute it to chemical variations in specific pathways contributing to the cell cycle, thus describing an advantageous phenotypic trait that does not originate from a specific genotype.

Given these considerations, the following *Assumptions* seem reasonable and will be used in the foregoing analysis:

1. There exist numbers *d*_+_ ≥ 0 and *r*_−_ *<* 0 *< r*_+_ such that *λ*_*i*_, *λ*_*h*_ : (*d*_+_, +∞) ⟶ (*r*_−_, *r*_+_).
2. The effective proliferation rates are strictly increasing as a function of the resource *c* and verify that *λ*_*i*_(*c*) *> λ*_*h*_(*c*) for every *c* ∈ (*d*_+_, +∞).
3. There holds that 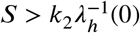.

#### Summary of results

Both models (1) and (2) share the same qualitative dynamical signature, which our numerical simulations confirm to be robust across a wide range of parameter values. Starting from a spatially localised initial condition, the system first undergoes a transient during which populations and resource adjust toward a local coexistence state. This is followed by the emergence of a coupled travelling wave pattern that propagates steadily through the domain: in the one-population case, the cell density and oxygen concentration move together at a common speed; in the two-population case, the fitter population invades while the host retreats, again at a common wavespeed, with the oxygen profile reorganising in concert.

The theoretical analysis of these travelling wave patterns is carried out on a quasi-steady-state (QSSA) reduction of the resource equation, which replaces the oxygen PDE by an algebraic constraint and reduces each model by one equation. Dimensional analysis identifies the small parameter *ϵ* = 1/(*k*_2_*T*) governing the validity of this approximation; numerically, the QSSA is found to be accurate over a parameter range considerably broader than the strict *ϵ* 0 1 regime. For the single-population reduced model, the structure is that of a monostable reaction-diffusion equation Fisher (1937); Kolmogorov et al. (1937); Hadeler and Rothe (1975), and the invasion speed is given by the following formula:

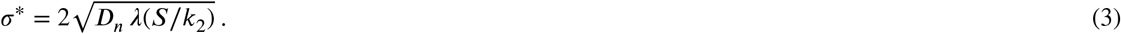

For the two-population reduced model, the analysis is more involved: the system does not have a planar phase portrait, but under a change of variables it can be recast as a monotone cooperative system Volpert et al. (1994), which yields existence of monotone coupled travelling waves. A rigorous lower bound on the invasion speed, together with a conjecture for the exact value given by

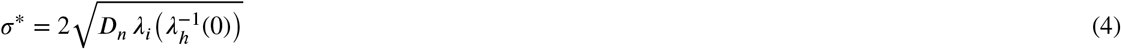

-compare with (3), are established and validated numerically both for the reduced model and, by extrapolation, for the full model over a wide parameter range. The analysis is detailed in Sections 2 and 3 respectively.

### 2. One species model: Tissue proliferation and travelling waves

As discussed in the introduction, solutions of (1) are expected to be well described, after an initial transient, by coupled oxygen–population travelling waves. By a coupled travelling wave we mean that both the population density and the oxygen concentration are given by profiles of constant shape moving at a common speed, i.e. solutions of the form *u*(*t, x*) = *u*(*x*– *σt*) = *u*(*ξ*) for some constant *σ >* 0. By convention we take *σ >* 0, so that in one spatial dimension the population proliferates from left to right and, in doing so, depletes the resource, whose profile likewise retreats to the right. Travelling wave solutions for (1) are therefore represented by real profiles *N* and *C*:

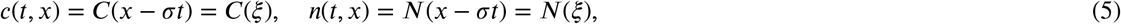

where the wave speed *σ* is the same for both unknowns.

#### Remark 2.1

*Let us describe in more detail a specific application of our abstract framework, as it is helpful to motivate why the general assumptions on the net proliferation rate λ. In the original application in de la Cruz et al. (2017), λ arises after coarse-graining the model with age structure. There λ is given, for each value of c, as the unique solution of*

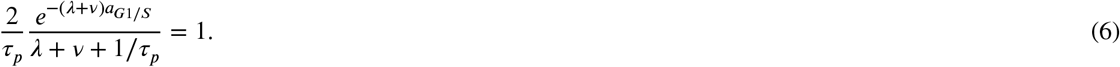

*Here a*_*G*1*/S*_ *is a function of c that captures some relevant features of a cell-cycle model -more specifically, it represents a cellular division age, whereas ν stands for a death rate and τ*_*p*_ *for the typical duration of a cell cycle. In this particular situation, it turns out that*

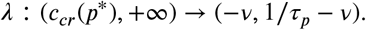

*The original model already assumes that ν <* 1*/τ*_*p*_, *so that the range of λ is consistent with our hypothesis. Moreover c*_*cr*_(*p*^*^) *is a strictly positive constant -representing a critical oxygen threshold, so the domain fits in our framework as well. Concerning the increasing character of λ, we notice that*

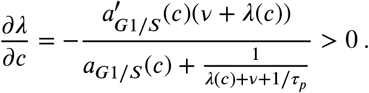

*Therefore, this specific model fits into our framework. Moreover, numerical simulations in de la Cruz et al. (2017) show that coupled oxygen-population waves are formed after an initial transient. See de la Cruz et al. (2017) for more details*.

Both the numerical simulations of (1) and the theoretical analysis of coupled travelling patterns are facilitated by a previous adimensionalisation of the model, a procedure that we describe next.

### 2.1. Adimensionalisation and numerical scheme

We introduce 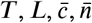 typical values for time, length, oxygen concentration and population concentration. In this way, we adimensionalise the unknowns via

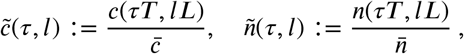

which, when evaluating the original equations (1) in points of the form (*τT, lL*), are seen to satisfy

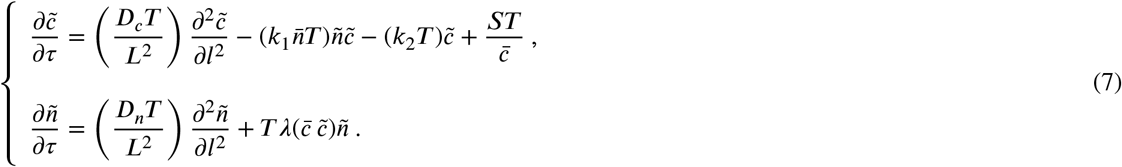

In what follows we choose

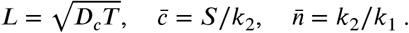

Therefore, we get

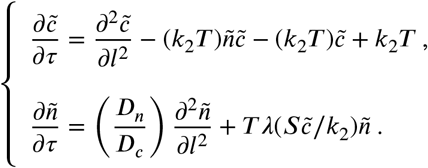

On letting *ϵ* := 1/(*k*_2_*T*) and dividing the first equation by *k*_2_*T* we arrive to

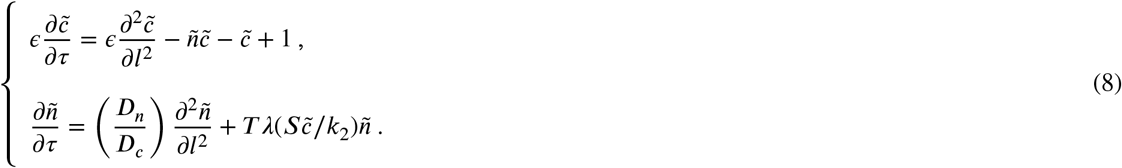

#### Remark 2.2

*For the specific model in de la Cruz et al. (2017) we set T* = *τ*_*p*_ *in what follows. Although this need not be the case for the general abstract models we consider, to think of T as representing the duration of a cell cycle will provide a convenient terminology in what follows*.

#### Equilibrium states and numerical initialisation

The coupled population–oxygen system admits a unique homogeneous equilibrium determined by the balance between oxygen supply, consumption, and the cellular proliferation threshold. In the quasi–steady limit for oxygen, the fast relaxation of the oxygen dynamics yields the algebraic relation

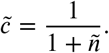

The non–trivial equilibrium is selected by the condition 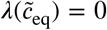, which, for the proliferation functions considered here, occurs at a critical oxygen level 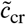. Consequently, the equilibrium population density is given by 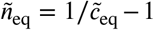 and does not depend on transport parameters such as diffusion ratios or oxygen rates. The equilibrium oxygen concentration observed in numerical simulations is found to coincide with the unique root of the implicit growth–rate equation *λ*(*c*) = 0, computed independently using the same cell–cycle model employed in the PDE system. All numerical simulations are initialised from this homogeneous equilibrium, introducing a localised spatial perturbation to trigger invasion. This choice ensures that simulations corresponding to different parameter values start from an identical tissue state, allowing a direct comparison of propagation dynamics. In particular, it enables a systematic assessment of the effects of the diffusion ratio *D*_*r*_ := *D*_*n*_/*D*_*c*_ and the oxygen relaxation parameter *ε* = 1/(*k*_2_*T*) on wave speed and profile, as well as a quantitative evaluation of deviations from the quasi–steady approximation arising from finite *ε* effects.

#### Spatial nondimensionalisation and numerical resolution

The spatial coordinate is nondimensionalised using the characteristic diffusion length

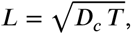

where *D*_*c*_ denotes the oxygen diffusion coefficient and *T* is the characteristic time scale of the system. Introducing the dimensionless spatial variable 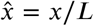, the spatial domain is defined as 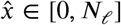, where *N*_*ℓ*_ represents the total number of characteristic diffusion length scales included in the computational domain. The domain is discretised into *N* spatial grid points, leading to a dimensionless spatial step

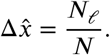

In dimensional variables, the spatial resolution is therefore given by

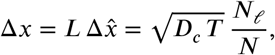

and the physical size of the domain corresponds to *N*_*ℓ*_*L*. This formulation allows the numerical resolution and the physical extent of the domain to be controlled independently: increasing *N* refines the spatial grid, while increasing *N*_*ℓ*_ enlarges the physical domain without affecting resolution.

*Conditions for resolving travelling wave fronts*. travelling wave solutions in reaction–diffusion systems are characterised by spatial gradients occurring over a length scale of order *L*. In order to numerically resolve such wave fronts, the spatial discretisation must satisfy

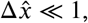

ensuring that several grid points lie within one characteristic diffusion length. In practice, reliable resolution of the wave profile and propagation speed requires 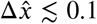, corresponding to at least ten grid points per diffusion length. In addition, the total domain size must be sufficiently large, *N*_*ℓ*_ ≫ 1, so that the travelling wave can fully develop and reach its asymptotic propagation regime before interacting with the domain boundaries. Together, these conditions imply the numerical requirement

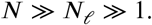

#### Dimensional and dimensionless wave speeds

Wave propagation speeds are naturally obtained in dimensionless form as 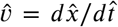, where 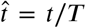 denotes the dimensionless time. The corresponding dimensional wave speed is recovered via the relation

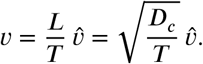

Equivalently, when the wave position is tracked numerically in terms of grid indices, the dimensional speed can be computed directly as

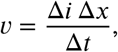

where Δ*i* is the displacement measured in grid points over a dimensional time interval Δ*t*. Provided the spatial discretisation satisfies the resolution criteria discussed above, the computed wave speed is independent of the grid spacing and reflects the intrinsic propagation dynamics of the model.

#### Temporal nondimensionalisation and time discretisation

Time is nondimensionalised using the characteristic duration of the cell cycle, denoted by *T* (= *τ*_*p*_), which provides a biologically meaningful reference scale for the dynamics. Introducing the dimensionless time variable 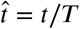, one unit of 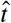 corresponds to one complete cell cycle. In the numerical implementation, each cell cycle is further subdivided into a prescribed number of subcycles, *n*_sub_, yielding a dimensionless time step

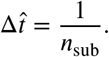

Accordingly, the dimensional time step is given by 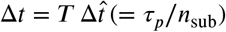.

#### Numerical scheme details

To simulate (8) we use a finite difference scheme. To advance in time each species we use a Crank–Nicolson method when dealing with the spatial diffusion, while the reaction terms are handled explicitly. To manage the species coupling we use a splitting technique. We work on a finite spatial interval with homogeneous Neumann boundary conditions. We use a uniform spatial grid, with *N* nodes. The method is unconditionally stable, and the time increment is taken as 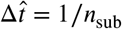, as specified above.

#### Emergence of travelling waves

A couple of representative examples are shown in Figure 1. Here we compare two different choices for the oxygen-dependent proliferation rate *λ*(*c*), using the same representative set of macroscopic parameters. In the first formulation, *λ*(*c*) is obtained from the coarse-grained cell-cycle model 6, where oxygen controls the G1/S transition time,

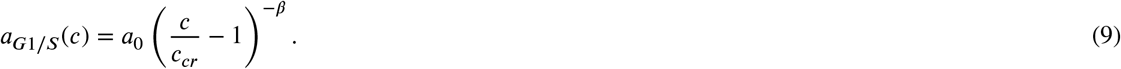

**Figure 1.**
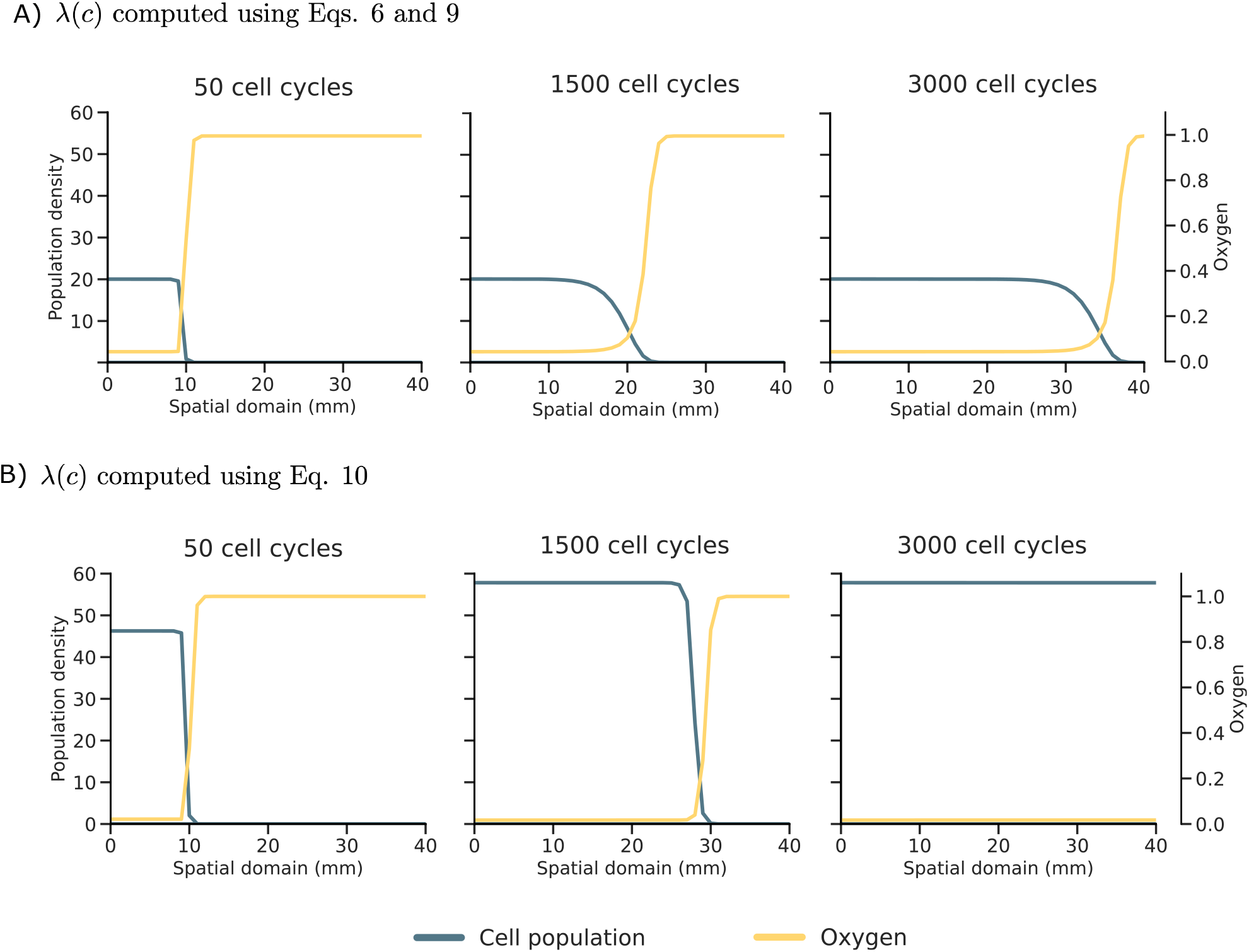
Simulations of (8) for two choices of the effective proliferation rate *λ*. (A) *λ* is computed using Eq. 6 and Eq. 9 from the cell-cycle coarse-graining model. *D*_*n*_ = 10^−*p*^ mm^2^/s, *D*_*c*_ = 10^−3^ mm^2^/s (*D*_*r*_ = 10^−4^), *k*_2_ = 1 s^−1^,*a*_0_ = 216591. The rest of the parameters can be found in 1. Starting from a localised initial condition, the population grows until it reaches its coexistence equilibrium with oxygen, after which a coupled travelling wave develops and advances toward the right. (B) *λ*(*c*) follows Eq. 10, where *α* = 6.7, rest of the parameters as in 1. Both the wave speed and profile shape differ between the two cases, but in both instances the coupled travelling wave structure emerges robustly after the initial transient.

Thus, the effective proliferation rate emerges indirectly from oxygen-dependent cell-cycle progression. In the second formulation, *λ*(*c*) is prescribed directly through the phenomenological response

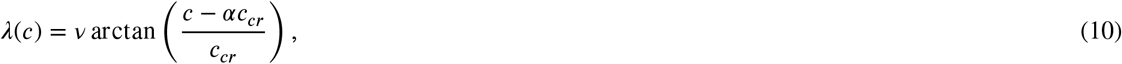

where *αc*_*cr*_ defines the oxygen level at which the net growth rate changes sign. The comparison between these two choices tests whether travelling-wave formation depends on the detailed microscopic form of cell-cycle regulation, or whether it is a robust consequence of oxygen-dependent effective fitness.

We can observe the transient dynamics and the evolution of the coupled travelling structure. We clearly see how the cell population and oxygen concentration initially relax toward a spatially uniform equilibrium over a wide spatial range, before the emergence of a coherent travelling wave, which subsequently governs the long-term invasion dynamics.

In what follows we explore this phenomenology systematically. Table 1 collects a representative set of biologically grounded parameter values, taken from the literature or callibrated for the cell-cycle model, which define the reference point for the parameter exploration below.

**Table 1.** Dimensional and dimensionless parameters used in the travelling-wave simulations. Cited values were taken from the literature, whereas uncited values correspond to model normalisations, calibrated cell-cycle parameters, or parameter ranges used for numerical exploration.

| Parameter | Value | Units |
| --- | --- | --- |
| $\tau_p$ | 4800 | s |
| $D_c$ | 0.001 (Ayensa-Jiménez et al., 2020) | m<sup>2</sup>/s |
| $D_n$ | $D_r D_c$ | m<sup>2</sup>/s |
| $D_r$ | $\{10^{-5}, 5 \times 10^{-5}, 10^{-4}, 5 \times 10^{-4}, 10^{-3}, 5 \times 10^{-3}\}$ | dimensionless |
| $k_2$ | $\{0.01, 0.05, 0.1, 0.5, 1, 5, 10\}$ | s<sup>-1</sup> |
| $k_1$ | 0.5 | density<sup>-1</sup>s<sup>-1</sup> |
| $S$ | 1 | mol s<sup>-1</sup> m<sup>-1</sup> |
| $v_h, v_i$ | $4.16667 \times 10^{-6}$ (de la Cruz et al., 2016) | s<sup>-1</sup> |
| $a_{0,h}, a_{0,i}$ | 216591, 130000 | s |
| $\alpha_h, \alpha_i$ | 6.7, 1.7 | dimensionless |
| $F_s$ | 1 | dimensionless |
| $c_{cr}$ | 0.01 (de la Cruz et al., 2016) | dimensionless |
| $\beta$ | 0.2 (de la Cruz et al., 2016) | dimensionless |

Let us discuss the parameter combinations that appear in (8). The parameter *T* can be thought of as the typical duration of the proliferative cell cycle; this is the meaning of *τ*_*p*_ in de la Cruz et al. (2017) -see Remark 2.2, so that ge *T* implies slow proliferation and vice versa. We fix *T* = *N*800 s throughout, which sets the biological timescale of tumour growth and allows us to focus attention on the remaining parameters. With *T* fixed, variations in *ϵ* = 1/(*k*_2_*T*) correspond directly to variations in *k*_2_, the rate of oxygen loss independent of cellular consumption.

#### On the robustness of the emergent dynamics

Whether a well-defined travelling wave front forms depends on the parameter regime. For large *D*_*r*_, cells disperse rapidly relative to oxygen transport and the wave structure does not emerge within the computational domain.

Throughout the simulations *D*_*c*_ and *τ*_*p*_ are kept fixed, so that the characteristic length scale 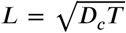 remains constant across all parameter explorations and the physical domain size does not change between runs.

Parameter variation is therefore performed through the diffusion ratio

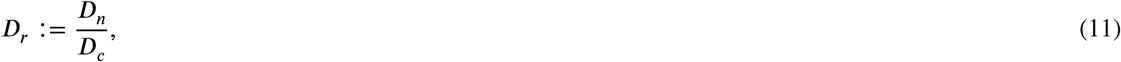

which controls cellular motility relative to oxygen diffusion. Since *D*_*c*_ is fixed, varying this ratio effectively corresponds to changing the cell diffusion coefficient *D*_*n*_ while preserving the underlying spatial scale set by oxygen transport.

This choice allows us to isolate the effect of relative cellular mobility on system dynamics without introducing changes in the geometry or physical size of the domain. In biological terms, the oxygen penetration length and tissue-scale transport properties remain fixed, while cellular motility is varied with respect to this reference scale.

### 2.2. Quasi-steady state approximation

To facilitate the analysis of (1) we can perform a quasi-steady state approximation (QSSA hereafter), taking for granted that the oxygen concentration reaches a locally homogeneous approximation very fast, i.e. the oxygen evolution equation would resemble

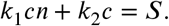

In other words, the approximation posits that

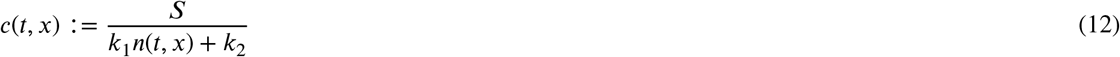

and this expression is subsequently substituted into the equation for *n*. Then the QSSA reduced model in physical units reads

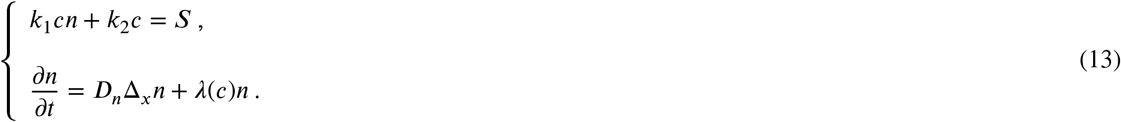

Having in mind the reformulation (8), we expect this approximation to work well when *ϵ* ≪ 1. Our numerical simulations show that this is also true for a broader range of parameters; we perform such simulations over the QSSA for (8), which reads

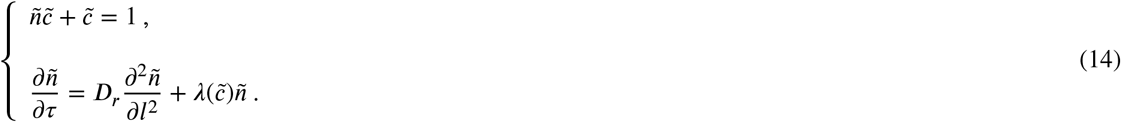

Here we introduced

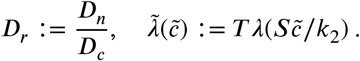

Thanks to the QSSA, the travelling wave problem for (14) becomes analytically tractable. The resulting wave-speed estimate is then extrapolated to the full model (1), yielding formula (3); its accuracy is assessed numerically in Section 2.4.

### 2.3. Travelling wave analysis for the reduced model

A travelling wave is a solution of constant profile moving at constant speed, i.e. of the form *u*(*t, x*) = *u*(*x*–*σt*) = *u*(*ξ*) with *σ >* 0. In our situation with (1), the waves connect the pair of homogeneous equilibria *E*_−∞_ and *E*_+∞_:

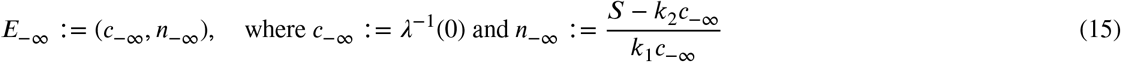

and

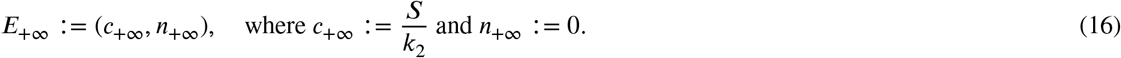

Note that these expressions make sense provided that Assumption 3 holds true.

Our numerical simulations (see e.g Figs. 1 and 2) show that both waves have the same speed. Hence we describe travelling wave solutions for our system as

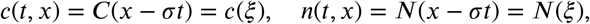

where the wave profiles now solve

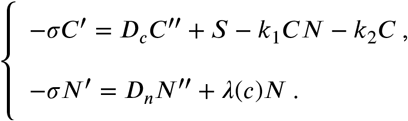

**Figure 2.**
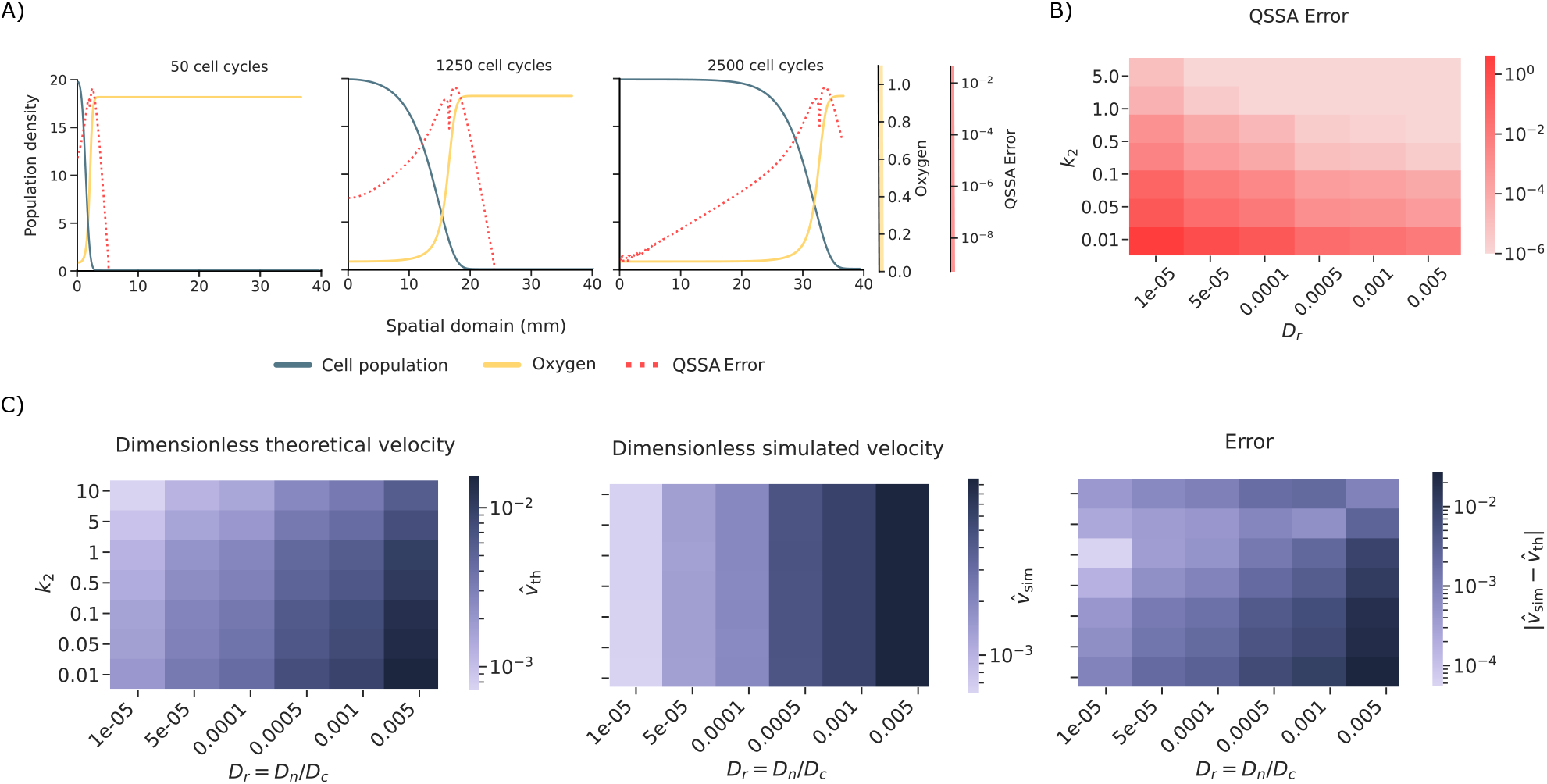
One-population travelling-wave dynamics and QSSA validation. (A) Spatial profiles of the cell population, oxygen concentration, and pointwise QSSA error at 50, 1050, and 1900 cell cycles. The effective growth rate is obtained from the oxygen-dependent transition time *a*_*G*1/*S*_ (*c*) = ._0_(*c*/*c*_*cr*_ − 1)^− *β*^ . The representative profiles in panel A correspond to *k*_2_ = 0.1 and *D*_*r*_ = *D*_*n*_/*D*_*c*_ = 0.0005; all remaining parameters are given in Table 1. The QSSA error remains small away from the front and is localised around the moving interface, where cell density and oxygen consumption vary most rapidly. (B) Error between the QSSA approximation (12) and the computational results across the parameter space (*D*_*r*_, *k*_2_), with *D*_*r*_ = *D*_*n*_/*D*_*c*_. Errors are larger for small *D*_*r*_, corresponding to sharper cellular fronts, and decrease for larger *k*_2_, consistent with faster oxygen relaxation. (C) Dimensionless theoretical velocity, simulated velocity, and absolute velocity error. The wave speed is mainly controlled by *D*_*r*_, while the simulated dependence on *k*_2_ is weak. This indicates that propagation is governed primarily by the effective fitness at the leading edge. Larger errors occur for faster waves, where finite-domain effects make the fitted velocity more sensitive to the available observation window.

When we perform the QSSA and pass to the travelling wave coordinates via (5), what we get is

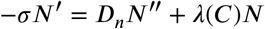

coupled with

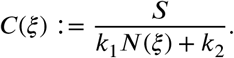

We complement this with the conditions

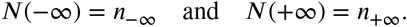

This simplified version allows for a fully analytical treatment.

Actually, it is convenient to work things out for the adimensional version (14), which allows for a systematic comparison with numerical simulations for the full model. Our travelling wave ansatz would be

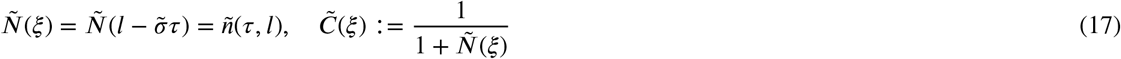

where 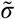 is the adimensional wave speed. The population profile satisfies

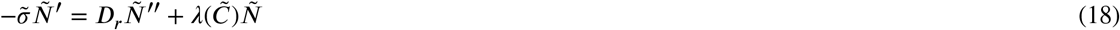

with limit conditions

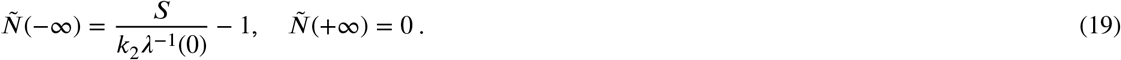

In this setting we can show the following (see the Supplementary Material for a full proof):

#### Proposition 2.3

*Assumptions 1–3 for the one-species model hold. Consider*

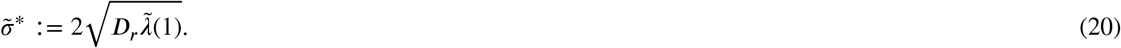

*Then, for each* 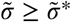, *there exists a monotonically decreasing travelling profile that solves* (17)*–*(18)*–*(19). *Actually*, 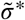 *is the minimal admissible travelling wave speed for this model*.

This result is readily translated to the dimensional version (13) and in turn it furnishes an accurate approximation of the dynamics for (1). In order to get the formula for the dimensional reduced model, note that the adimensional wave speed can be rewritten as

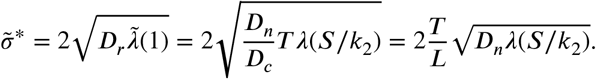

Hence, returning to the original units, we recover formula (3) for the observable wavespeed *σ*^*^. Consistent with the classical theory of monostable reaction-diffusion equations Hadeler and Rothe (1975), invasion occurs at the linear (minimal) speed *σ*^*^. Although this prediction is derived for the reduced model, the agreement with the wave speed measured in numerical simulations of the full model (1) is quantitatively good, as we show next.

### 2.4 Numerical assessment of the QSSA

The spatial structure of the QSSA error is examined in Figure 2A for the one-species model. The error remains small and spatially localised near the invasion front, while being negligible in the bulk regions behind and ahead of the wave. Figure 2B provides a global assessment of the QSSA error over the parameter space (*D*_*r*_, *k*_2_) explored in this work. Across the parameter space, the QSSA error is larger for smaller values of *D*_*r*_. Since *D*_*r*_ = *D*_*n*_/*D*_*c*_, this corresponds to a regime in which cellular diffusion is small compared with oxygen diffusion, where the cellular profile is less spatially smoothed and the invasion front is sharper. The error also displays a clear dependence on the oxygen decay rate *k*_2_. Larger values of *k*_2_ correspond to faster oxygen relaxation, and the QSSA error decreases in this regime. This is consistent with the quasi-steady assumption: the QSSA is expected to be more accurate when oxygen equilibrates rapidly compared with the timescale of cellular motion and proliferation. Thus, Figure 2B provides a computational validation of the timescale-separation argument underlying the QSSA.

The wave speed is estimated from numerical simulations by tracking the position of maximum spatial gradient in the population density across a sequence of simulation snapshots, fitting a linear regression to the resulting front-position time series, and extracting the slope as the propagation speed. We display the errors incurred over a representative range of parameters in Figure 2 C. This shows the theoretical dependency on the oxygen decay parameter however this is not captured in the computational experiments, showing the fact that the oxygen relaxes fast and does not plays a role in the velocity of the wave. The error plot shows that the theoretical prediction (3) is accurate to within a few percent across the parameter ranges considered, with the error growing modestly as *D*_*r*_ increases toward the regime where front formation becomes marginal. However, the simulated velocity shows a much weaker dependence on this parameter and varies predominantly with *D*_*r*_. This indicates that, in the full simulations, the oxygen level ahead of the front is effectively buffered close to its equilibrium value, or that the growth function is already close to saturation in the relevant oxygen range. Thus, changing *k*_2_ strongly affects the oxygen relaxation dynamics and the QSSA error, but only weakly changes the leading-edge growth rate that determines the observed wave speed.

In this formulation of the model, the wave propagates into an initially empty and oxygenated region. The theoretical velocity is therefore controlled by the effective growth rate at the leading edge and by the cell diffusion coefficient. In agreement with this prediction, both theoretical and simulated velocities increase with *D*_*r*_, reflecting the relation *D*_*n*_ = *D*_*r*_*D*_*c*_. The absolute velocity error is larger in the faster-wave regimes, especially at high *D*_*r*_.

This increase should be interpreted with caution, as it does not necessarily indicate a breakdown of the travelling-wave approximation. Rather, larger values of *D*_*r*_ generate faster and more diffuse fronts: the position of the leading edge is less sharply defined, fewer data points are available for a robust linear regression before the wave reaches the domain boundary, and the diffuse front expands the spatial region over which the local oxygen environment regulates growth. The observed increase in absolute error therefore reflects a combination of front-detection uncertainty, finite-domain effects, and the comparison of larger propagation velocities, rather than a failure of the travelling-wave approximation itself. Note that this is distinct from the pointwise QSSA error discussed above: sharper fronts can increase the local QSSA error, since oxygen consumption varies rapidly across the interface and diffusive smoothing becomes locally important, yet the same sharpness can make the wave speed easier to estimate, as the front position and leading-edge oxygen level are then more clearly defined.

## 3. Two species model: competition and the role of phenotypic traits

We use the model (2) to describe the competition of two populations which are spatially segregated. Those populations differ in their respective values of a phenotypic trait, say *p*^*^, that in turn controls the proliferation rates depending on oxygen consumption. In the case of the model described in Remark 2.1, the phenotypic trait that controls the proliferation rate is ._0_, which represents the age at which a cell reaches the G1/S transition in the cell cycle under optimal conditions. Beyond this point, cells become capable of dividing, provided that sufficient oxygen is available. In a resourced mediated domain this age is oxygen dependent and it is calculated as in Eq. 9. where *c* is the oxygen concentration and *c*_*cr*_ the critical oxygen value above the which cells can divide. In this two species model, the value of *a*_0_ differs between the two populations and therefore determines their distinct proliferation dynamics. Specifically, *a*_G1/S_ is used to compute the net birth–death rate (Eq. 6), so different values of 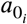 and 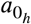 will lead to different growth *λ*_*i*_ and *λ*_*h*_, respectively for the invader and the host populations.

An alternative way of encoding phenotypic differences is to prescribe the oxygen-dependent net growth rate directly, using the one formulated in Eq. 10, where *α* shifts the effective oxygen threshold of the population. In this formulation, phenotypic differences are represented by different values of *α*: a smaller *α* corresponds to a population able to sustain growth at lower oxygen concentrations, whereas a larger *α* represents a population requiring higher oxygen availability. Unlike the cell-cycle based formulation, this expression does not derive the growth rate from the G1/S transition age, but provides a smooth phenomenological approximation of oxygen-dependent proliferation.

The parameter *α* modulates the oxygen sensitivity of a population. Indeed, the argument of the arctangent can be written as *c*/*c*_*cr*_ − *α*, so changing *α* shifts the growth-response curve along the oxygen axis. The value *c* = *αc*_*cr*_ represents the effective oxygen level at which the transition between unfavourable and favourable growth occurs. Therefore, smaller values of *α* correspond to phenotypes that can sustain growth under lower oxygen availability, whereas larger values of *α* correspond to phenotypes requiring higher oxygen levels to proliferate efficiently. In a two-population setting, choosing *α*_*i*_ *< α*_*h*_ gives the invading population a growth advantage in oxygen-poor regions, since its net growth rate remains higher than that of the host over a broader range of resource concentrations.

To perform numerical studies of the competition dynamics, we divide the spatial domain in two parts and we populate them with a spatially homogeneous cell distribution with different proliferative phenotypes. Then we let it evolve. What we expect to see is that in a first stage the populations will readjust themselves to the local oxygen conditions. After a transient, the fittest population will gradually take over the other, seemingly as a population wave with roughly constant speed. Those effects are actually described by solutions to (2). In order to fulfill this program, let us explain first how to adimensionalise the model.

### 3.1. Adimensionalisation and the QSSA for the competition model

Here we proceed as in Section 2.1. Let us denote by 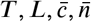 typical values for time, length, oxygen concentration and population concentration. We introduce

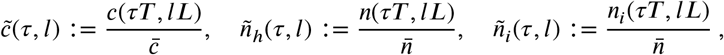

where, as before, we take

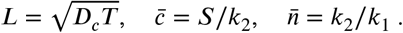

In this way we rewrite (2) as

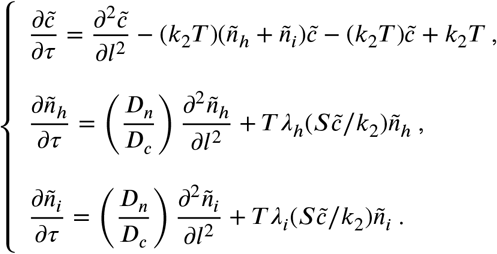

Now we introduce

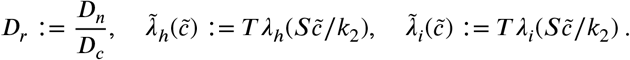

We also let *ϵ* := 1/(*k*_2_*T*) and divide the first equation by *k*_2_*T* . As a result, we arrive at

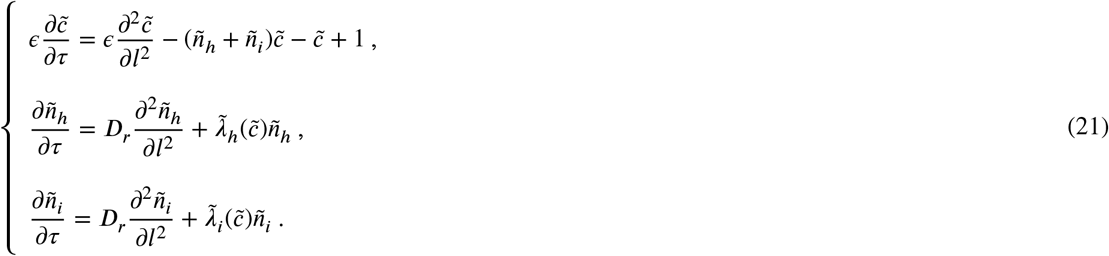

Two representative scenarios were simulated to obtain Figure 3. We see that in both cases we have a transient recalibration dynamics, after which the aforementioned takeover dynamics sets in. The non-transient behavior can be corroborated by analysing a simplified model, which we shall do below.

**Figure 3.**
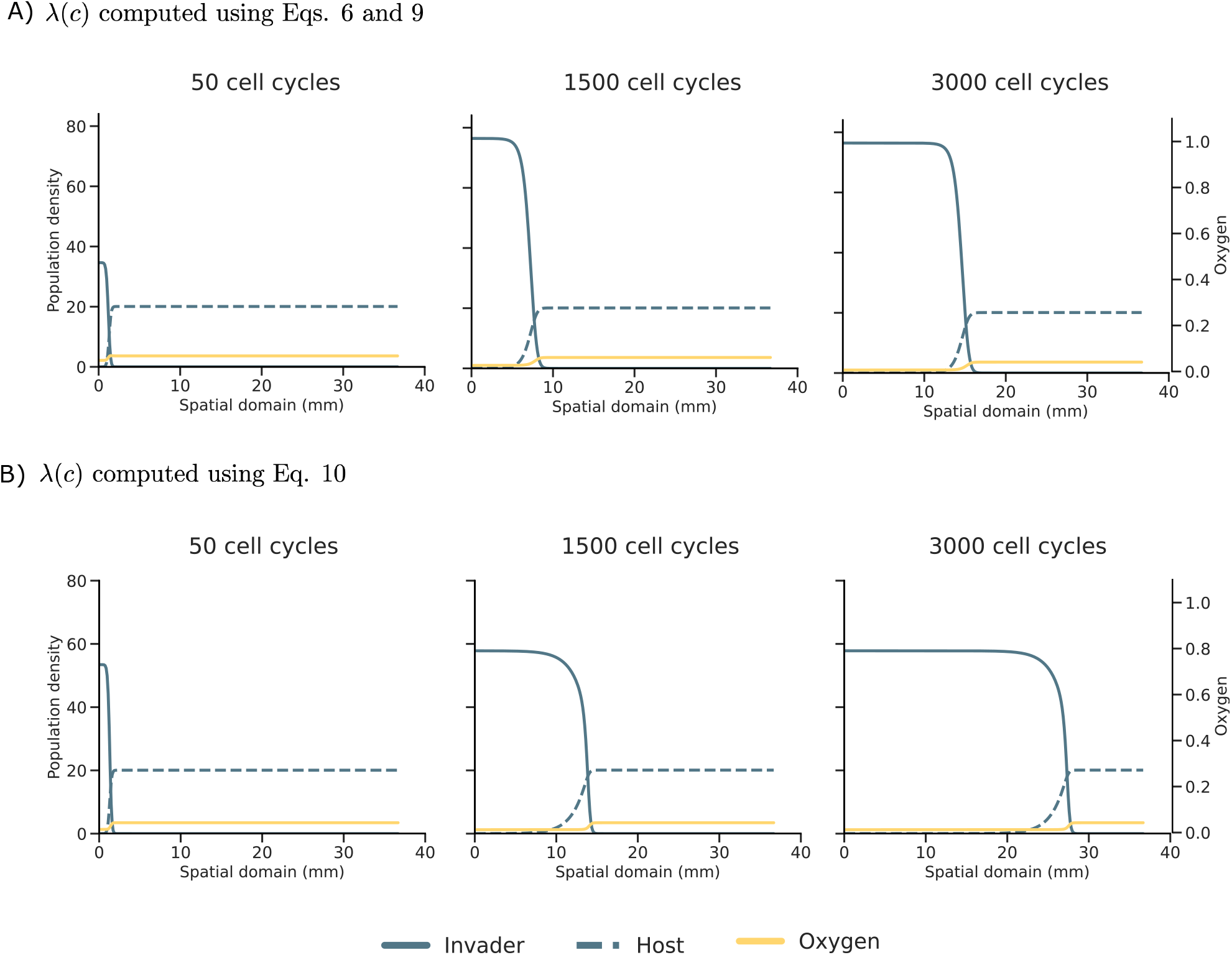
Dynamics of the competition model (21). Travelling-wave dynamics for two alternative oxygen-dependent growth laws in the two-population competition model. Snapshots are shown at 50, 1500 and 3000 cell cycles. The simulations shown correspond to *k*_2_ = 1.0 and *D*_*r*_ = *D*_*n*_/*D*_*c*_ = 10^−4^; all remaining parameters are given in Table 1. Solid blue curves denote the invader population, dashed blue curves denote the host population, and yellow curves denote the oxygen concentration, plotted on the secondary vertical axis. (A) Competition dynamics for the threshold-based cell-cycle model (Eq. 6), where the oxygen-dependent transition time is given by *a*_,1/*S,J*_ (*c*) = *a*_0,*J*_ (*c*/*c*_*cr*_ − 1)^− β^, *J* = *i, h* (Eq. 9). The invader has a lower cell-cycle delay, *a*_0,*i*_ *< a*_0,*h*_, and therefore a higher effective fitness under the same oxygen conditions. (B) Competition dynamics for the arctangent growth model, where *λ*_*J*_ (*c*) = arctan((*c* − *α*_*J*_ *c*_*cr*_)/*c*_*cr*_), *J* = *i, h* (Eq. 10). Here the invader has a lower oxygen sensitivity threshold, *α*_*i*_ *< α*_*h*_, allowing it to maintain positive growth in regions where the host is growth-limited. In both cases, the invader progressively replaces the host through a travelling invasion front, illustrating how differences in oxygen-dependent fitness can drive spatial competitive exclusion.

We might simplify the model (21) by assuming the oxygen to be in a locally homogeneous equilibrium. The QSSA for (2) thus amounts to substitute the first equation in (2) by

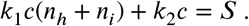

When we perform this approximation in the nondimensional version (21), what we are assuming is that

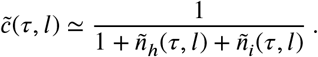

Therefore, we will study the following simplified, nondimensional model:

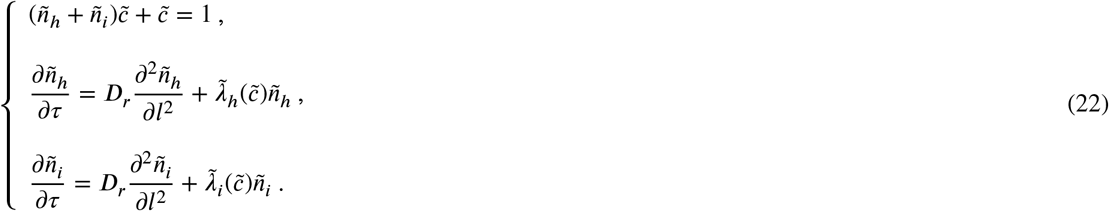

As in the case of (14), we expect this approximation to (21) to work fine at least for *ϵ* ≪ 1.

### 3.2. Travelling wave analysis

First we introduce specific notations, similarly to what we did in Section 2.3. We start with the dimensional version (2). Let us describe travelling wave solutions as

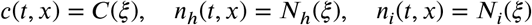

with *ξ* = *x* – *σt*. By convention, let us take *σ >* 0 and the invasion taking place from left to right. That is, *N*_*i*_ (*ξ*) is a decreasing profile and *N*_*h*_(*ξ*) is an increasing profile, should they exist. Actually, the wave profiles would solve

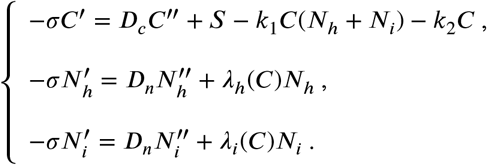

However, when we introduce the QSSA we reduce the problem to the last two equations above, where we substitute

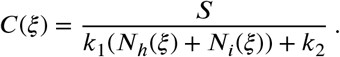

This is a four-dimensional, non-autonomous dynamical system; simpler than the starting six-dimensional one, yet not straightforward to manage.

Let us address now the boundary conditions. The homogeneous equilibria (*c*(−∞), *n*_*h*_(−∞), *n*_*i*_(−∞)) and (*c*(+∞), *n*_*h*_(+∞), *n*_*i*_(+∞)) associated with this situation are the following:

- At *ξ* = −∞ we expect *n*_*h*_ to vanish, *n*_*h*_(−∞) = 0. Hence we expect

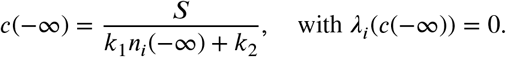

In this way

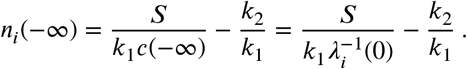

- At *ξ* = +∞ we expect *n*_*i*_ to vanish, *n*_*i*_(+∞) = 0. Hence we expect

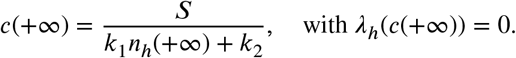

In this way

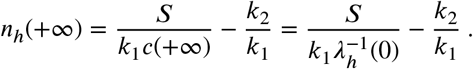

When we reformulate the problem with respect to the nondimensional version (22), we use the ansatz

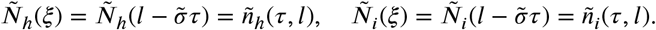

Here 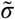 is the adimensional wave speed. As per the QSSA, we take

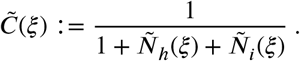

The wave profiles for both populations would satisfy

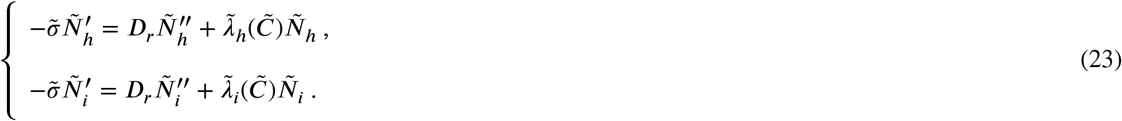

In this adimensional formulation, the boundary conditions read

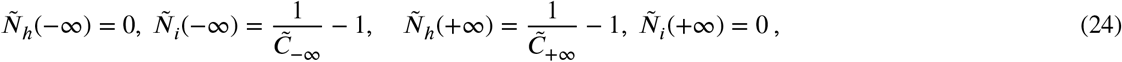

where

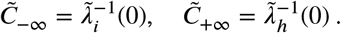

We may now show the following (proofs are given in the Supplementary Material):

#### Proposition 3.1

*Assumptions 1–3 for the two-species model hold true. Then, there is some* 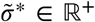, *such that the following properties hold: For each* 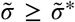, *there exists a pair of travelling profiles* (*Ñ*_*h*_(_*ξ*_), *Ñ*_*i*_(*ξ*)) *that solves* (23), *together with the boundary conditions at* _*ξ*_ = ±∞ *specified in* (24). *Each component Ñ*_*h*_, *Ñ*_*i*_ *is monotone, with N*_*h*_ *increasing and N*_*i*_ *decreasing. Furthermore*, 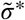 *is bounded from below by the linear speed*,

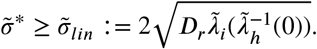

To obtain the wavespeed formula for the model with physical units, note that

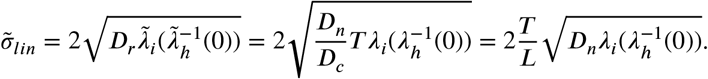

In this way we recover formula (4) for the wavespeed 1 of the reduced model in physical units, which we can use to approximate that of the original competition model (2).

### 3.3. Numerical study of the emergent competition pattern

The validity of the QSSA was assessed by comparing the oxygen profile obtained from the full reaction–diffusion system with the corresponding quasi-steady approximation. Representative spatial profiles are shown in Figure 4A. As in the single-population case, the pointwise QSSA error remains small over most of the spatial domain and is mainly concentrated around the propagating invasion front. Away from the front, the error is typically very small, often reaching values below 10^−6^ whereas localised peaks appear near the cellular interface. This localization is expected: in the bulk regions, cell densities vary slowly and the oxygen field is close to local equilibrium, while near the invasion front the cellular density, and therefore the oxygen consumption term, changes rapidly over a short spatial scale.

**Figure 4.**
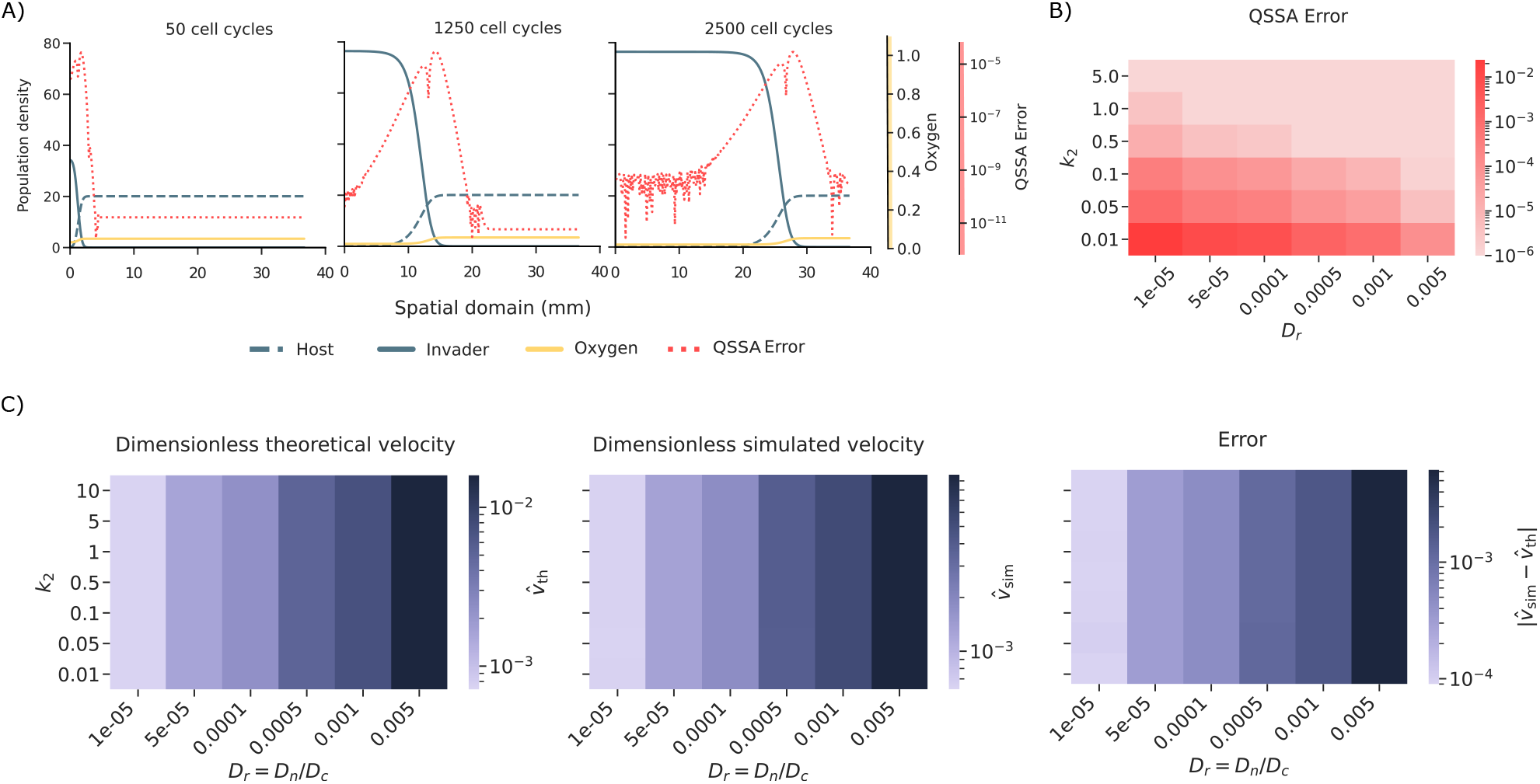
Two-population travelling-wave dynamics and QSSA-based speed approximation using the threshold-based cell-cycle growth law. The effective growth rates are obtained from the oxygen-dependent transition times *a*_*G*1/*S,J*_ (*c*) = *a*_0,*J*_ (*c*/*c*_*cr*_ − 1)^− β^, *J* = *i, h*, with distinct parameters for the host and invader populations. *D*_*r*_ = 0.0005, *k*_2_ = 0.1 and the rest of parameters are indicated in Table 1. (A) Representative spatial profiles of the host population, invader population, oxygen concentration, and pointwise QSSA error at 50, 1250, and 2500 cell cycles. The invader progressively replaces the host through a travelling competition front, while the oxygen field adapts to the moving cellular interface. The QSSA error is mainly localised around the front, where cell densities and oxygen consumption vary most sharply, and remains small in the bulk regions. (B) Global QSSA error across the explored parameter space (*D*_*r*_, *k*_2_), where *D*_*r*_ = *D*_*n*_/*D*_*c*_ is the ratio between cell and oxygen diffusion coefficients, and *k*_2_ is the oxygen decay rate. The error is larger for smaller *D*_*r*_, corresponding to sharper cellular interfaces, and decreases as *k*_2_ increases, consistent with faster oxygen relaxation improving the quasi-steady approximation. (C) Dimensionless theoretical velocity, simulated velocity, and absolute velocity error for the invasion front. The theoretical speed is computed from the invader effective growth rate evaluated at the oxygen level set by the resident population, while the simulated speed is obtained by tracking the moving competition front. Both theoretical and simulated speeds are primarily controlled by *D*_*r*_, whereas the dependence on *k*_2_ is weak. The largest absolute errors occur in the fastest-wave regimes, where finite-domain effects make numerical speed estimation more challenging and fronts are broader.

A systematic quantification over the explored parameter space is reported in Figure 4B. The QSSA error decreases as oxygen relaxation towards the quasi-equilibrium becomes faster, for instance for larger values of the oxygen decay rate *k*_2_, and remains substantially smaller outside the front region than at the front itself, analogously to the single population model. This confirms that the main limitation of the local QSSA is not the description of the bulk oxygen field, but rather the narrow transition layer where the moving cellular interface produces sharp spatial gradients in oxygen consumption. In particular, sharper fronts can generate larger local peaks in the QSSA error because the algebraic QSSA neglects the diffusive, smoothing term, *D*_*c*_*∂*_*xx*_*c*. Thus, the QSSA should not be interpreted as a uniformly accurate pointwise approximation across the whole domain, but as an effective reduced description whose accuracy is high away from steep transition regions.

The impact of this approximation on the travelling-wave description is summarised in Figure 4C. The theoretical and simulated wave speeds display the same qualitative dependence on the diffusion ratio *D*_*r*_ = *D*_*n*_/*D*_*c*_, and the corresponding absolute velocity error remains controlled across most of the explored parameter space. This indicates that the travelling-wave estimate can remain accurate even when the QSSA exhibits localised discrepancies near the front. This is because the wave speed is primarily determined by the effective oxygen-dependent growth rate at the leading edge, rather than by pointwise agreement of the oxygen field everywhere in the domain.

Throughout the parameter exploration, *D*_*c*_ and *T* (= *τ*_*p*_) were kept fixed, while the cell diffusion coefficient was varied through the ratio 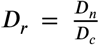 This choice isolates the effect of relative cellular mobility while preserving the oxygen transport scale and the physical size of the computational domain. For each simulation, the front position was extracted from a sequence of output snapshots. The front was identified either from the relevant population profile or, in the two-population case, from the competition interface. Only frames for which the front remained inside the spatial domain were retained. This procedure provides a robust estimate of the propagation speed while filtering out small frame-to-frame numerical fluctuations.

There are two sources of uncertainty in this measurement. First, for large cellular diffusion, the wave can travel rapidly across the domain, leaving fewer data points available to fit the wave speed before the front reaches the boundary. Second, as the front becomes broader, its precise position is less sharply defined and can depend on the front-detection criterion. These effects are a numerical limitation of finite-domain travelling-wave simulations, rather than a failure of the analytical approximation itself. For this reason, the number of selected frames and the quality of the linear fit were monitored together with the velocity error.

The velocity approximation shows different but consistent behaviour in the single-population and two-population settings. In the single-population case, the wave propagates into an initially empty and oxygenated region. The theoretical velocity is therefore controlled by the cell diffusion coefficient and by the oxygen-dependent growth rate at the leading edge. Both the theoretical and simulated velocities increase with *D*_*r*_, reflecting the increase of *D*_*n*_ = *D*_*r*_*D*_*c*_. However, the simulated velocity shows only a weak dependence on *k*_2_. This suggests that, in the full simulations, changes in *k*_2_ affect the oxygen relaxation dynamics and the QSSA error, but do not strongly alter the effective growth rate experienced by cells at the leading edge. In other words, the wave speed is controlled primarily by the effective fitness at the front, rather than by the far-field oxygen balance alone. This behaviour is illustrated in Figure 2C for the single-population simulations.

In the two-population case, the propagating structure is not an expansion into empty space, but a competitive invasion front in which one population replaces the other. The theoretical speed is therefore computed from the invader growth rate evaluated at the oxygen level for which the resident population has zero net growth, rather than from the far-field oxygen concentration *S*/*k*_2_. The relevant quantity is thus the effective growth rate of the invader in the microenvironment set by the resident population. Despite this additional coupling, the approximation captures the main trend of the simulated speeds across the explored values of *D*_*r*_. Both theoretical and simulated velocities vary primarily along the *D*_*r*_ direction, while the dependence on *k*_2_ is weak. This supports the interpretation that, in the competitive setting, the speed is mainly controlled by cellular mobility and by the oxygen-dependent fitness difference between the resident and invading populations.

Across both modelling settings, the largest absolute velocity errors occur in the fastest-wave regimes. This is partly a finite-domain effect: faster waves reach the boundary sooner, reducing the number of reliable frames available for the linear regression used to estimate *D*_sim_. In addition, larger *D*_*r*_ produces broader and faster fronts, for which the location of the leading edge is less sharply defined. The velocity error should therefore be interpreted together with the number of selected frames and the quality of the front-position fit.

Overall, Figures 4A–C support the use of the QSSA as an effective reduced description of the oxygen field and show that the travelling-wave approximation captures the main propagation behaviour of the full simulations. The QSSA is most accurate in the bulk, while its main pointwise limitation is localised near sharp invasion fronts. Nevertheless, these localised discrepancies do not prevent accurate prediction of the dominant wave-speed trends, because propagation is governed primarily by the effective oxygen-dependent fitness experienced at the invasion edge.

## 4. Discussion and conclusions

In this paper, we have studied the emergence of coupled travelling wave patterns in a family of coarse-grained reaction-diffusion models of resource-mediated tissue competition. The central modelling insight is that phenotypic heterogeneity at the level of the cell cycle — encoded here through an effective, resource-dependent proliferation rate — is sufficient to confer a replicative advantage to one cellular lineage over another, without any requirement for genetic mutation or differential access to the resource itself. Even minor differences in proliferation efficiency can be amplified, through the spatial feedback between local cell density and local resource availability, into a winner-takes-all invasion dynamic operating at the tissue scale. This mechanism provides a complement to genetic variability as an explanation for the emergence of dominant phenotypic subpopulations, in line with observations in related modelling frameworks Chisholm et al. (2015); Ardaševa et al. (2020).

The dynamics we observe unfold in two stages. First, during a transient phase, the competing populations readjust their local densities in response to the available resource, reaching a spatial coexistence state in which the fittest population is present at a higher proportion. In the second phase, this coexistence state propagates through the medium as a coupled travelling wave: the invading population advances, the host population retreats, and the resource concentration reorganises in concert, all at a common wavespeed. This two-stage sequence — transient readjustment followed by coherent wave propagation — is a robust feature of the model across a wide range of parameter values, and mirrors the kind of progressive competitive displacement observed in tissue competition settings Gatenby and Gawlinski (1996); Martínez-González et al. (2012).

A key contribution of this work is to show that the quasi-steady-state approximation (QSSA) for the resource dynamics provides a quantitatively accurate description of this behaviour across a broad and biologically relevant parameter regime. The QSSA reduces the coupled population–resource system to a simpler form that retains the dominant feedback structure while becoming tractable to rigorous analysis. For the single-population model, the reduced system is of Fisher–KPP type Fisher (1937); Kolmogorov et al. (1937); Hadeler and Rothe (1975), and the invasion speed is determined by the linearisation about the unstable equilibrium in the wake of the wave. For the two-population competition model, the reduced system is a four-dimensional non-autonomous dynamical system. The analysis of coupled travelling waves in such systems is notoriously difficult, and most existing results in the literature rely on numerical simulation Mascia et al. (2021); Moschetta and Simeoni (2019); Lorenzi et al. (2025a) or perturbative and singular-perturbation approaches Burie et al. (2006); Crossley et al. (2023); El-Hachem et al. (2021); Marchant et al. (2001, 2000); Sherratt (1993); Tyson and Keener (1988). Analytical existence results are considerably rarer, and typically require the imposition of specific structural assumptions that allow the use of tools such as the Conley index Conley and Gardner (1984); Mischaikow and Reineck (1993), topological degree Gardner (1982), shooting arguments Dunbar (1984); Gallay and Mascia (2022); Huang (2016); Carrillo et al. (2025); Colson et al. (2021), or fixed-point theorems Zhang et al. (2016). In the present work, we exploit the fact that, under a suitable change of variables, the reduced two-population system can be recast as a monotone cooperative system Volpert et al. (1994). This structure yields existence of coupled travelling wave solutions with monotone profiles, and provides a rigorous lower bound on the invasion speed. Our numerical simulations confirm that this lower bound is in practice achieved, and that the resulting wave-speed estimate agrees well with the speed observed in the full model.

The wave-speed estimate has a clear biological interpretation: it is strongly dominated by the proliferation rate of the fittest population evaluated at the critical resource concentration below which the host population can no longer sustain positive net growth. This is a manifestation of the more general principle that invasion speed in monostable reaction-diffusion systems is set by the growth properties of the advancing population in the region ahead of the front, where the resource is most abundant. The fact that the host population’s properties enter only through this threshold concentration reflects the asymmetry of the competition: the invader’s fitness at the resource level that the host can just tolerate is the decisive quantity.

The numerical simulations show that the dominant dependence of the wave speed is on the relative cellular mobility, through *D*_*n*_ = *D*_*r*_*D*_*c*_. In both the single-population and two-population settings, simulated speeds increase with *D*_*r*_, in agreement with the travelling-wave scaling. By contrast, the dependence of the simulated speed on the oxygen decay rate *k*_2_ is weak, despite the fact that *k*_2_ affects the oxygen relaxation profile and the pointwise QSSA error. This indicates that *k*_2_ does not control the invasion speed directly in the explored regime. Instead, its effect is filtered through the oxygen-dependent fitness experienced at the leading edge of the wave. In the single-population case, this corresponds to the effective proliferation rate of cells at the front. In the two-population case, it corresponds to the growth rate of the invader in the oxygen environment set by the resident population. This supports the view that the relevant biological control parameter is not the full oxygen profile, nor the resource decay rate in isolation, but the effective fitness landscape generated by the coupling between oxygen availability and cell-cycle progression.

Several directions remain open and merit further investigation. First, the asymptotic stability of the travelling fronts obtained here has not been established; this is a technically demanding problem for higher-dimensional systems, but would considerably strengthen the predictive value of the wave-speed estimates. Second, the perturbative relationship between the full model and its QSSA reduction could be made rigorous, for instance through slow-manifold theory in the spirit of Fenichel’s theorem, which would provide sharper quantitative control on the parameter regimes in which the approximation is valid. Third, the present analysis is restricted to one spatial dimension and to at most two competing populations; extensions to higher spatial dimensions and to systems with more than two lineages would be of direct biological interest, although the monotonicity structure exploited here would be lost in the latter case, making analytical progress substantially harder. Finally, it would be valuable to connect these deterministic predictions to stochastic and individual-based descriptions of the same competition process de la Cruz et al. (2017), in order to assess the robustness of the travelling wave phenomenology to demographic fluctuations and to understand the corrections imposed by finite cell-number effects.

In summary, this work demonstrates that resource-mediated competition, operating through phenotypic heterogeneity at the cell-cycle level and without genetic variability, is sufficient to generate robust coupled travelling wave invasion patterns in spatially extended tissue models. The quasi-steady-state reduction provides both an accurate numerical approximation and an analytically tractable framework, and the resulting wave-speed estimates are robust across a wide range of macroscopic parameters. The dominant determinant of invasion speed is the proliferation capacity of the fittest lineage at the resource threshold of its competitor; a simple, measurable, and biologically interpretable quantity.

## Acknowledgment

N.B. and P.G. have been supported by Grant PID2022-141802NB-I00 (BASIC) and the research network RED2022-134573-T funded by MCIN/AEI/10.13039/501100011033 and, by ‘ERDF A way of making Europe’. J. C. has been partially supported by Grant C-EXP-265-UGR23 funded by Consejería de Universidad, Investigación e Innovación and ERDF/EU Andalusia Program; by Grant PID2022-137228OB-I00 funded by the Spanish Ministerio de Ciencia, Innovación y Universidades, MICIU/AEI/10.13039/501100011033 and ‘ERDF/EU A way of making Europe’; and by Grant QUAL21-011 (Modeling Nature) Consejería de Universidad, Investigacion e Innovacion of the Junta de Andalucía. R.O.-B. acknowledges support from the Program Horizon Europe Marie Sklodowska-Curie Post-Doctoral Fellowship (HORIZON-MSCA-2023-PF-01). Grant agreement: 101149877. Project acronym: NFROGS. Part of this work was carried out during the three-months secondment period of R.O.-B. at the Departamento de Matemática Aplicada, Universidad de Granada, which he thanks for their hospitality. J. C. also acknowledges partial support from project NFROGS and thanks Dipartimento di Informatica, Università di Verona for their hospitality; part of this work was carried there while visiting R.O.-B. The work of T.A. has been funded by the AEI grant PID2024-162434OB-I00 “ERDF A way of making Europe”. Further support has been provided by the Spanish Research Agency (AEI), through the Severo Ochoa and Maria de Maeztu Program for Centres and Units of Excellence in R&D (CEX2020-001084-M). T.A. thanks the CERCA Programme/Generalitat de Catalunya for institutional support.

## Declaration of generative AI and AI-assisted technologies in the manuscript preparation process

During the preparation of this work, the author(s) used ChatGPT (OpenAI) in order to assist with copy-editing tasks, including improving the clarity and phrasing of prose in the main text, consolidating redundant passages, and checking internal consistency of notation and terminology across sections. After using this tool, the author(s) reviewed and edited the content as needed and take full responsibility for the content of the published article.

## 1 Supplementary Material

### 1.1 On the one-species model

Here we supply a proof for Proposition 2.3 in the main text. This is concerned with the equation

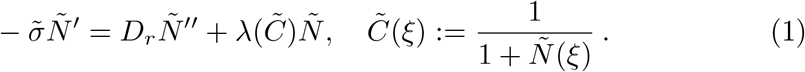

We rewrite it as

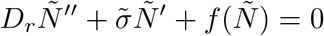

where

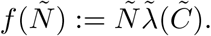

Notice that *f* is positive on (0, *ñ*_−∞_) and vanishes on *Ñ* = 0 and *Ñ* = *ñ*_−∞_. Moreover,

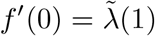

and one checks that *f*^*′*^ (*ñ*_−∞_) < 0. Therefore, *f* is a standard monostable nonlinearity. It is then known that there exists a minimal admissible speed 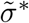 and travelling fronts from *ñ*_*-1*_ to 0 for (1) exist if and only if 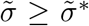. Using a classical result by Hadeler and Rothe [4], we show that invasion occurs at the linear speed -i.e. the speed of propagation for small perturbations about the stable equilibrium point, that is

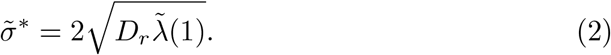

Indeed, as shown in [4, Corollary 9] one has

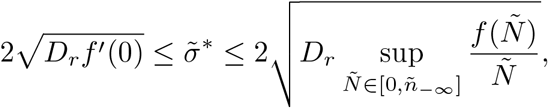

that is

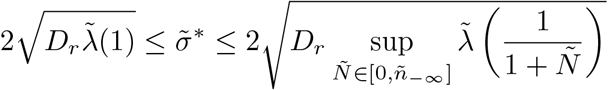

and since 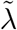 is strictly increasing, (2) follows. The fact that the profiles of the fronts are decreasing is well known, see for instance [1, Theorem 4.2].

### 1.2 On the two-species model

Here we provide a proof for Proposition 3.1 in the main text. This is concerned with the two-species model

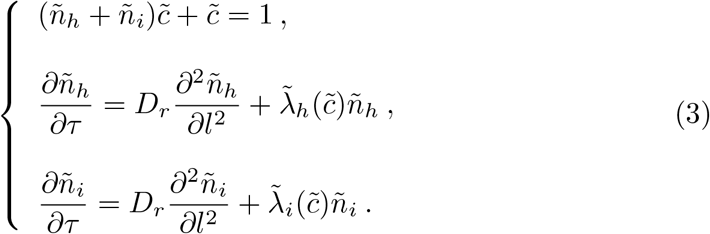

We split the argumentation in two different parts.

#### Step 1: Reformulation of the simplified competition model as a monotone system

In the absence of diffusion, the system (3) writes as

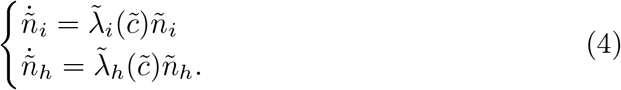

Writing *u*_1_ = *ñ*_*i*_ and *u*_2_ = *ñ*_*h*_, the system (4) becomes

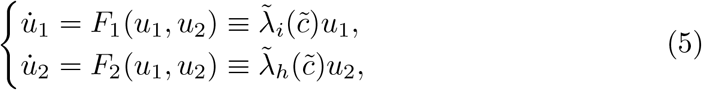

where

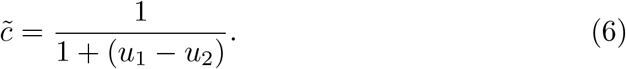

We restrict our attention to the region of interest, which is [0, *ñ*_*i*_(− ∞)] × [−*ñ*_*h*_(+ ∞), 0]. From (6) we obtain

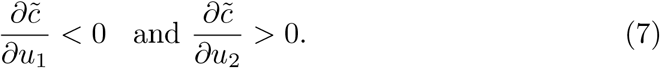

The equilibrium states of the system (5) are

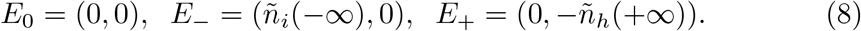

The system (5) is monotone. Indeed, by using (7) we obtain

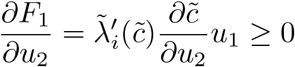

and

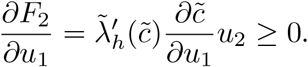

Let us now compute the eigenvalues of the system (5) linearized at each of its equilibrium points, listed in (8). For an arbitrary point, the differential of *F* = (*F*_1_, *F*_2_) reads

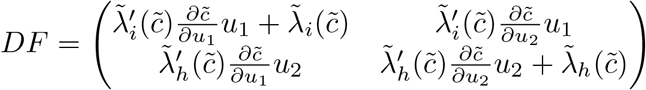

Hence,

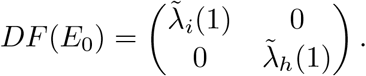

Thus, the eigenvalues at *E*_0_ are 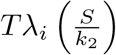 and 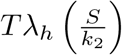, which are both positive due to Assumption *3* for the two-species scenario -i.e. 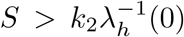. As a consequence, *E*_0_ is a repeller.

Let us compute now

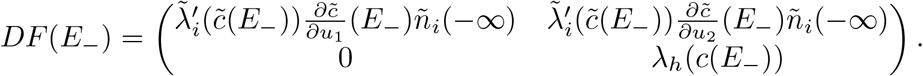

One eigenvalue is 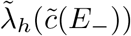, which is smaller than 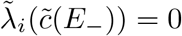, and hence is negative. The other eigenvalue is 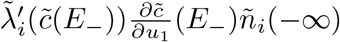, which is negative. That is, *E*_−_ is an attractor.

Finally, we compute

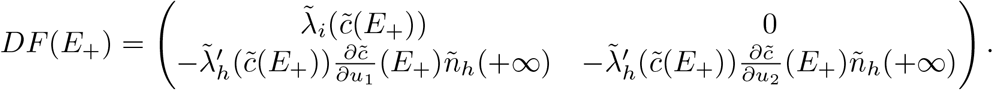

One eigenvalue is 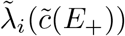, which is larger than 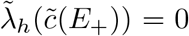, and hence is positive. The other eigenvalue is 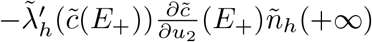, which is negative. In summary, *E*_−_ is a saddle point.

Therefore, we can apply [5, Chapter 3, Theorem 4.2 and Remark below] and find that there exists 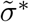 (given by the variational formula (11) below) such that for all 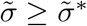 there exists a travelling wave solution for (3) at speed 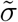 which is monotone in each component and goes from *E*_−_ to *E*_+_ -moreover, for 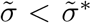 there are no such solutions. The monotonicity property for the profiles is clearly visible in the numerical simulations below and has an appealing interpretation -note that the invasion profile decreases in the invading direction, while the profile for the retreating population has the opposite trend, and both profiles are intertwined, hence their sum is roughly constant.

#### Step 2: Lower bound for the wave speed

Here we work with the coupled, nondimensional profile equations, that is,

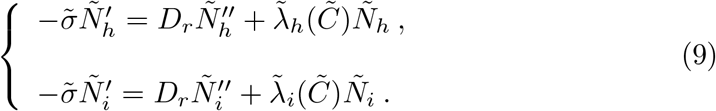

This is reformulated as a first order system in the usual way:

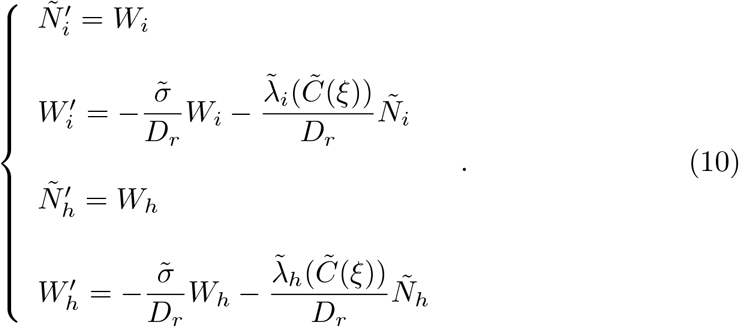

Recall that the existence of coupled travelling waves is equivalent to the existence of heteroclinic connections between *P*_−∞_ and *P*_+∞_, where we denote *P*_− ∞_ = (*n*_*i*_(∞ −), 0, 0, 0) and *P*_+∞_ = (0, 0, *n*_*h*_(+∞), 0). The linearization at the arrival point *P*_+∞_ reads

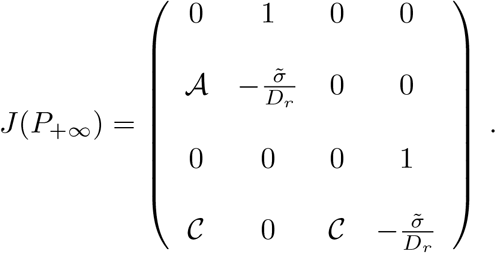

Here we abridged

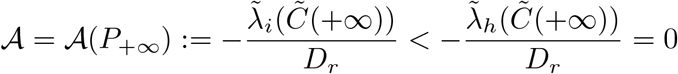

and

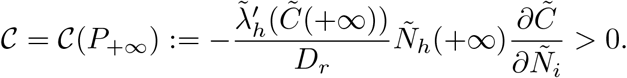

The associated eigenvalues read:

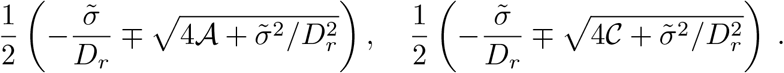

The last two eigenvalues are clearly real (one positive and one negative). The first two will be real (in which case they are both negative) provided that the following condition holds:

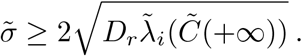

In case this condition does not hold, we would get two eigenvalues which are complex conjugate, one real positive eigenvalue and one real negative eigenvalue.

Likewise, we can also compute the associated eigenvectors:

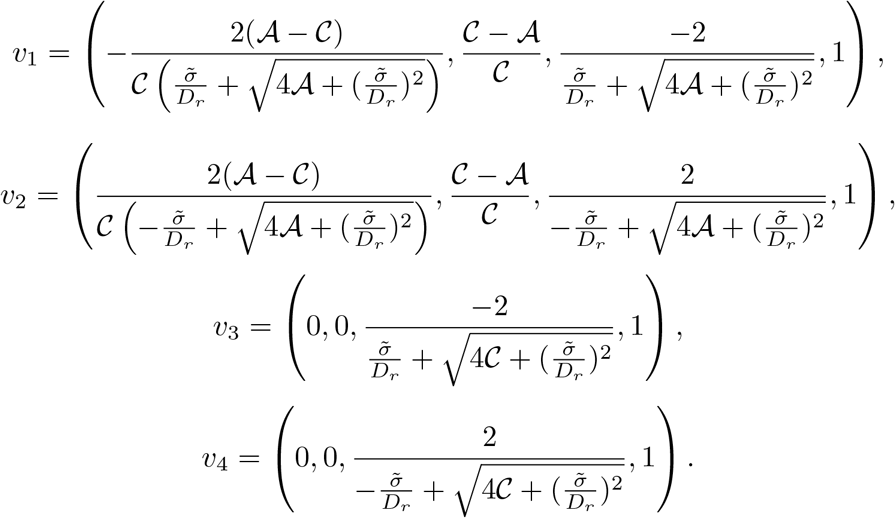

The eigenvectors associated to the complex eigenvalues have no zero components. Specifically, the third component is clearly nonvanishing, which implies that the travelling profiles thus obtained would violate the nonnegativity condition. Therefore, the former lower bound on 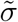 is a necessary condition for the existence of coupled travelling waves as defined in the main text.

##### Remark 1.1 (On upper bounds for the wave speed)

*The variational method introduced in [5] allows in principle to retrieve the invasion wavespeed for the coupled system. Indeed, the former theorem gives also a variational formula for the wave speed:*

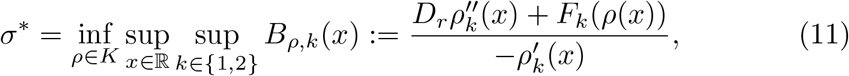

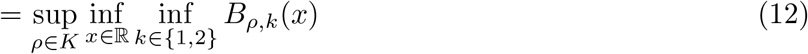

*Equation* (12) *is provided in [5, Chapter 5, Theorem 7*.*1] and holds under some extra technical assumptions. The set K is defined as the set containing the functions ρ* ∈ *C*^2^(ℝ, ℝ^2^) *such that* lim_*x*→*±*∞_ *ρ*(*x*) = E_*±*_ *and* 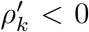 *for k* ∈ *{*1, 2*}. As pointed out elsewhere, this mechanism is notoriously difficult to put to good use (formally taking ρ as the wave profiles themselves gives a tautological statement; the point is to be able to approximate those within K in such a way that the given variational formula yields nontrivial information). We have not been able to extract useful information by means of this device; any improvements along this line would be most wellcome*.

### 1.3 On parameter values

#### Diffusion ratio

The ratio *D*_*r*_ := *D*_*n*_/*D*_*c*_ controls the relative spatial scales of cell movement and oxygen transport. In most biologically relevant regimes this ratio is small, reflecting the fact that oxygen diffuses considerably faster than cells migrate; in this limit, cell dynamics is governed primarily by growth rather than diffusion. Varying *D*_*r*_ allows us to explore the transition from a reaction-dominated to a transport-dominated regime.

Based on values reported in the literature [2], *D*_*r*_ can span several orders of magnitude. Combining the smallest reported cell diffusion coefficient, *D*_*n*_ = 6.6 × 10^−12^ cm^2^/s, with the largest reported oxygen diffusion coefficient, *D*_*c*_ = 2.0 × 10^−5^ cm^2^/s, yields

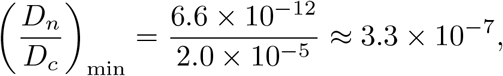

while the converse extremes give

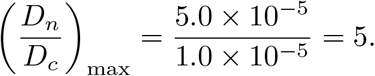

In practice, our exploration range for *D*_*r*_ is further constrained by numerical considerations. For *D*_*r*_ ≳ 0.01, cells move sufficiently rapidly that a well-defined travelling wave front does not form within the spatial domain considered, regardless of the domain size tested. Consequently, not all values of *D*_*r*_ in the biologically admissible range permit a systematic travelling wave analysis. Interestingly, the QSSA introduced below remains reasonably accurate even in these parameter regimes where no front is formed (see Figures 2 and 4).

#### Decay constants

At those regions where the population for the full one-species model reaches a nontrivial equilibrium, the equilibrium values are

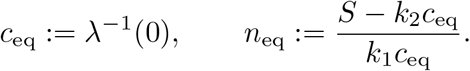

Hypothesis *3* -i.e. *S* > *k*_2_ *λ*^−1^(0)-guarantees that *n*_eq_ ≥ 0. Since *c*_eq_ is held fixed while *k*_2_ varies, the admissibility constraint *n*_eq_ ≥ 0 translates to the bound

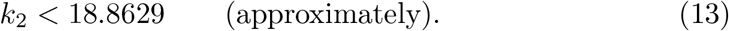

In nondimensional variables this equilibrium relation reads

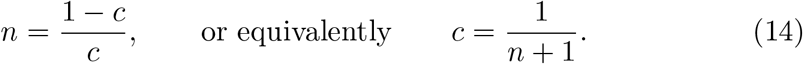

All simulations are initialised from the homogeneous equilibrium, with the nondimensional population density set to

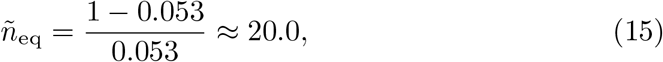

where the value 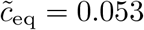 corresponds to the unique zero of λ for the cell-cycle model of [3]. This uniform initialisation ensures that all simulations are directly comparable across the parameter range explored.

##### Remark 1.2

*For the specific proliferation model in [3], the equilibrium oxygen concentration in the presence of cells is given by*

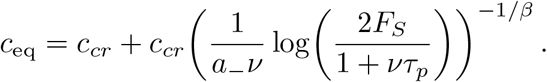

